# Mild Mitochondrial Impairment Activates Overlapping Longevity Pathways Converging on the Flavin-Containing Monooxygenase FMO-2

**DOI:** 10.64898/2026.02.10.705198

**Authors:** Jeremy M Van Raamsdonk

**Affiliations:** Department of Neurology and Neurosurgery, McGill University, Montreal, Quebec, Canada; Metabolic Disorders and Complications Program, and Brain Repair and Integrative Neuroscience Program, Research Institute of the McGill University Health Centre, Montreal, Quebec, Canada; Brain Repair and Integrative Neuroscience Program, Research Institute of the McGill University Health Centre, Montreal, Quebec, Canada; Division of Experimental Medicine, Department of Medicine, McGill University, Montreal, Quebec, Canada

**Keywords:** aging, lifespan, *C. elegans*, *fmo-2*, genetics, mitochondria, genetics

## Abstract

A mild impairment of mitochondrial function activates the hypoxia inducible factor (HIF-1)-mediated hypoxia stress response pathway leading to a HIF-1-dependent increase in lifespan. Lifespan extension resulting from HIF-1 stabilization is dependent on activation of flavin-containing monooxygenase-2 (FMO-2). In this work, we explored the role of *fmo-2* in the long lifespan of genetic mitochondrial mutants in *C. elegans*. We found that *fmo-2*, but not other *fmo* genes, are specifically upregulated in the long-lived mitochondrial mutants *clk-1, isp-1* and *nuo-6*. Disruption of *fmo-2* through RNA interference or genetic mutation shortens the lifespan of these mitochondrial mutants indicating that *fmo-2* is required for lifespan extension in these worms. Moreover, signaling molecules that have been shown to be involved in upregulation of *fmo-2* are also required for the long life of *clk-1, isp-1* and *nuo-6* mutants including HLH-30, NHR-49 and MDT-15. Finally, we examined the effect of multiple lifespan-promoting pathways in *clk-1* mutants on the expression of *fmo-2*. We found that in all cases, genes required for *clk-1* longevity are also required for the upregulation of *fmo-2* in *clk-1* worms. These genes included DAF-16, PMK-1, SKN-1, CEH-23, AAK-2, HIF-1 and ELT-2. Combined, this work advances our understanding of the molecular mechanisms contributing to longevity in the long-lived mitochondrial mutants and identifies FMO-2 as a common downstream effector of multiple pathways that modulate longevity.

## Introduction

Activation of the HIF-1 hypoxia pathway extends lifespan in *C. elegans* ^1, 2^. This can be achieved through either overexpression of *hif-1* or through disruption of *vhl-1*, which encodes the E3 ubiquitin ligase von Hippel-Lindau protein (VHL-1). Moreover, it has been shown that the HIF-1 hypoxia pathway is required for the extended longevity of multiple long-lived mitochondrial mutants including *clk-1, isp-1* and *nuo-6* ^3, 4^.

To gain insight into the mechanisms by which the HIF-1 hypoxia pathway promotes longevity, an RNAi screen was performed to identify RNAi clones that inhibit the decrease in age-associated autofluorescence in long-lived *vhl-1* mutants ^5^. They found that RNAi targeting the flavin-containing monooxygenase-2 gene *fmo-2* not only inhibits the reduction of autofluorescence in *vhl-1* worms but also decreases their extended longevity. Moreover, ubiquitously overexpressing *fmo-2* was found to be sufficient to increase lifespan. The lifespan extension in *fmo-2* overexpression worms was found to be independent of the stress-responsive transcription factors HIF-1, DAF-16 and SKN-1. Disruption of *fmo-2* was found to also prevent lifespan extension resulting from dietary restriction but not *daf-2* RNAi or *isp-1* RNAi ^5^.

In exploring mechanisms leading to *fmo-2* upregulation, it was found that the TFEB transcription factor HLH-30, which has been shown to regulate autophagy, is required for the full induction of *fmo-2* in response to hypoxia or dietary restriction ^5^. HLH-30 is also required for lifespan extension resulting from disruption of *vhl-1* or dietary restriction ^5, 6^, thereby providing further links between *fmo-2* expression and longevity.

A subsequent study showed that RNAi knockdown of transaldolase (*tald-1)* increases lifespan and activates *fmo-2* in a HLH-30- and PMK-1-dependent manner ^7^. Importantly, *fmo-2* is required for *tald-1* RNAi to increase lifespan as *tald-1* RNAi fails to extend longevity in *fmo-2* mutants. A further study into the mechanisms of *fmo-2* activation found that MDT-15, which encodes a subunit of the Mediator transcriptional coactivator complex, and NHR-49, which encodes a nuclear hormone receptor that acts with MDT-15 to regulate various aspects of metabolism, are both required for *fmo-2* activation ^8^. This study also demonstrated that the effect of NHR-49 on *fmo-2* activation is independent of HLH-30, and that SEK-1 and PMK-1 from the p38-meidated innate immune signaling pathway are also required for full *fmo-2* activation.^8^

Finally, it was shown that exposing worms to heat-inactivated, exopolysaccharide-producing strains of lactobacilli results in increased expression of *fmo-2* and extended longevity ^9^. The increase in lifespan was found to be dependent on *fmo-2* as well as *hlh-30* and *nhr-49*, which are involved in *fmo-2* activation ^9^.

In this work, we further explore the role of *fmo-2* in longevity. We find that *fmo-2* is upregulated in a specific group of long-lived mutant strains, which includes the long-lived mitochondrial mutants *clk-1, isp-1* and *nuo-6*. We find that FMO-2 and proteins involved in regulating FMO-2 expression, including HLH-30, SEK-1, PMK-1, NHR-49 and MDT-15 are all required for the long-lifespan of the mitochondrial mutants. Moreover, genetic pathways that affect longevity in *clk-1* mutants tend to also affect the expression of *fmo-2*. Overall, this work shows that *fmo-2* is important in mediating the lifespan extension in long-lived mitochondrial mutants. While this work provides additional support for the positive relationship between *fmo-2* expression and longevity, it also provides counter examples in which long-lived mutants (*eat-2, osm-5*) have extended longevity despite decreased expression of *fmo-2*.

## Materials and Methods

### Strains

Strains were maintained on NGM plates at 20°C and fed OP50 *E. coli* bacteria. The following strains were used in this study: N2(wild-type), *sod-2(ok1030), clk-1(qm30), isp-1(qm150), nuo-6(qm200), daf-2(e1370), glp-1(e2141), eat-2(ad1116), osm-5(p813), ife-2(ok306), fmo-2(ok2147), pmk-1(km25), ceh-23(ms23), aak-2(ok524)* and *hif-1(ia4)*. All strains were outcrossed a minimum of 3 generations. Some of these strains were crossed together to generate the following double mutant strains: JVR453 *clk-1(qm30);fmo-2(ok2147)*, JVR441 *nuo-6(qm200);fmo-2(ok2147)*, JVR227 *clk-1(qm30);pmk-1(km25)*, JVR099 *clk-1(qm30);ceh-23(ms23)*, JVR038 *clk-1(qm30);aak-2(ok524)*, and MQ1758 *clk-1(qm30);hif-1(ia4)*.

### RNA isolation

RNA was isolated from pre-fertile young adult worms as described previously ^10^. Briefly, worms from an overnight limited lay were collected and washed three times in M9 buffer before being frozen in Trizol. RNA was isolated from three biological replicates collected on different days.

### RNA sequencing

RNA sequencing was performed and processed previously ^4, 11-13^. Raw sequencing data is available on NCBI GEO: GSE179825, GSE93724 and GSE110984. An online tool for looking up genes of interest in this panel of long-lived mutants may be found here: **https://vanraamsdonk.shinyapps.io/mutant_comparison_viewer/**.

### Quantitative RT-PCR

For quantitative RT-PCR (qPCR), RNA was converted to cDNA using a High-Capacity cDNA Reverse Transcription Kit (Life Technologies) according to the manufacturer’s instructions. qPCR was performed using a FastStart Universal SYBR Green kit (Roche) in an AP Biosystems RT-PCR machine.

### Lifespan

For lifespan assays, pre-fertile young adult worms were picked from maintenance plates to lifespan plates containing 50 µM 5-fluoro-2’-deoxyuridine (FUdR). This concentration of FUdR completely prevents progeny from developing to adulthood but can affect the lifespan of specific strains ^14^. Throughout the lifespan, worms were checked every 2-3 days for survival by observation and gently prodding worms that are not moving voluntarily. Three biological replicates started on three separate days were performed.

### RNA interference

RNA interference (RNAi) was performed on plates containing 3 mM IPTG and 50 µg/ml carbenicillin. For lifespan experiments, worms were initially grown on RNAi plates with no FUdR. Once worms reached adulthood, they were transferred to RNAi plates containing 50 µM FUdR.

### Statistical Analysis

For lifespan studies, genotypes or RNAi clones were coded so that the identity of the strain or clones was not known by the experimenter. Graphpad PRISM Version 9.4.1 was used to prepare graphs and perform statistical analyses. Error bars indicate standard error of the mean. *p<0.05, **p<0.01, ***p<0.001, ****p<0.0001.

## Results

### *fmo-2* expression is increased in long-lived mitochondrial mutants

Previous work has demonstrated an increase in *fmo-2* expression in long-lived *vhl-1* mutants and under dietary restriction, and that in both cases *fmo-2* is required to extend longevity ^5^. In addition, overexpression of *fmo-2* is sufficient to increase lifespan ^5^. Given the role of *fmo-2* in lifespan extension in these different contexts, we sought to determine the extent to which *fmo-2* also plays a role in the extended longevity of the long-lived mitochondrial mutants: *clk-1, isp-1* and *nuo-6* ^15-17^.

As a first step, we examined the expression of *fmo-2* in the long-lived mitochondrial mutants. To do this, we utilized RNA sequencing data from a recent study in which we compared genes expression across a panel of nine long-lived mutants ^18^. In addition to the three long-lived mitochondrial mutants, this panel contained worms with decreased insulin/IGF-1 signaling (*daf-2*)^19^, worms with dietary restriction (*eat-*2)^20^, worms with decreased translation (*ife-*2)^21^, worms with decreased chemosensation (*osm-*5) ^22^, worms with germline ablation (*glp-1*) ^23^ and worms with elevated mitochondrial superoxide (*sod-2*) ^24^. Interestingly, we previously found that these nine long-lived mutants cluster into three distinct groups based on gene expression. The long-lived mitochondrial mutants are part of longevity group 1 which also contains *daf-2, glp-1* and *sod-2* worms, while *eat-2* and *osm-5* worms make up longevity group 2.

In examining the expression of *fmo-2* across these nine long-lived mutants, we found that *fmo-2* mRNA is significantly increased in all six of the group 1 long-lived mutants (*sod-2, clk-1, isp-1, nuo-6, daf-2, glp-1*), which includes the long-lived mitochondrial mutants (**Figure 1A**). The observation of increased *fmo-2* expression in *glp-1* worms is consistent with a previous study showing an increase in *glp-1* worms using an *fmo-2p::GFP* reporter strain ^8^. In contrast, *fmo-2* expression is significantly decreased in the group 2 longevity mutants *eat-2* and *osm-5* and unaffected in the group 3 *ife-2* mutants (**Figure 1A**). These findings suggest that *fmo-2* does not contribute to the extended longevity of *eat-2, osm-5* or *ife-2* mutants.

**Figure 1.**
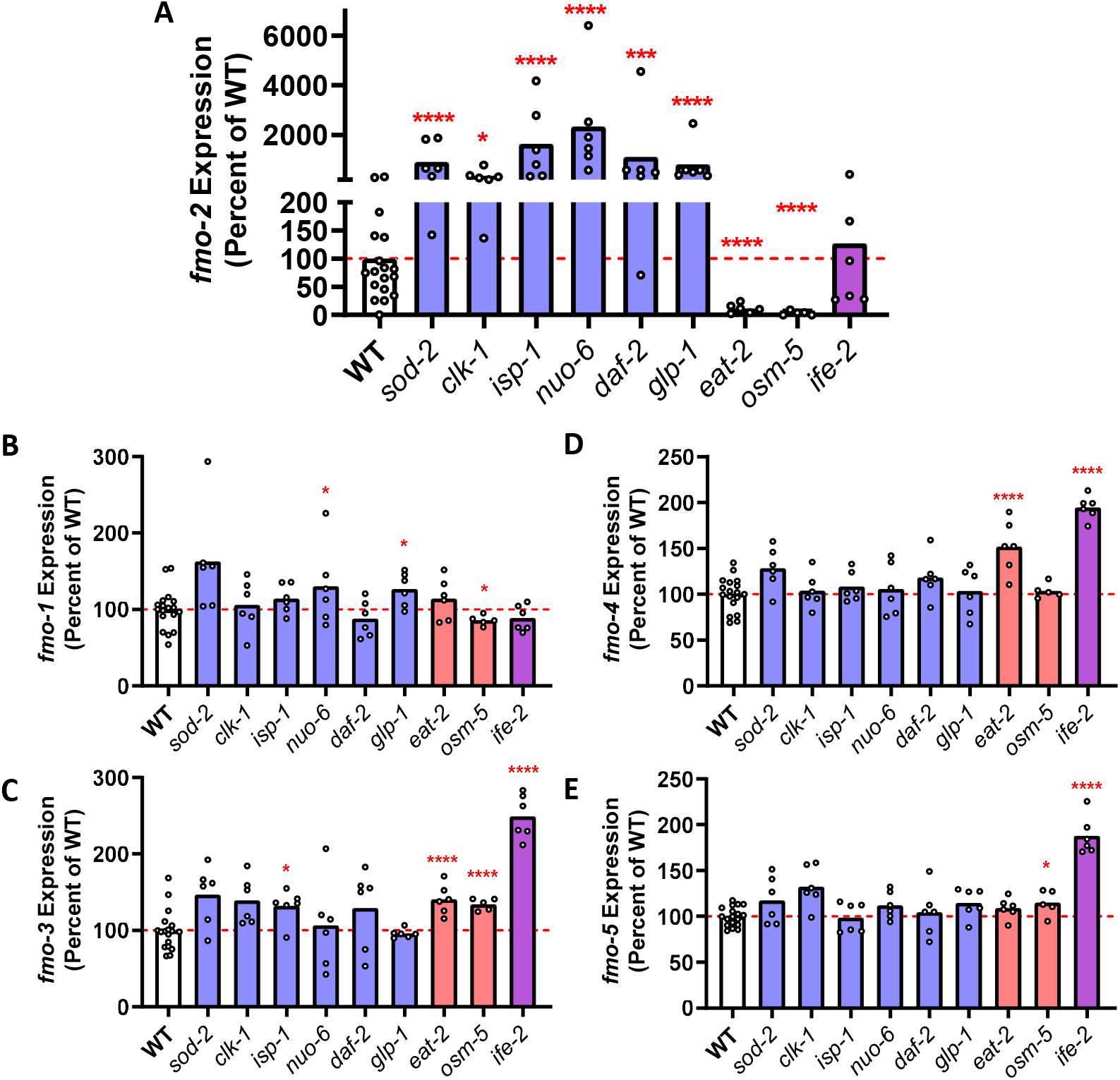
Expression of *fmo-2* mRNA is increased in long-lived mutants. (**A**) Long-lived mutants from longevity group 1 (*sod-2, clk-1, isp-1, nuo-6, daf-2, glp-1*) exhibit a significant upregulation of *fmo-2*, while long-lived mutants from longevity group 2 (*eat-2, osm-5*) show significantly decreased expression of *fmo-2. fmo-2* expression is unchanged in the group 3 longevity mutant *ife-2*. In contrast, expression of *fmo-1* (**B**), *fmo-3* (**C**), *fmo-4* (**D**) and *fmo-5* (**E**) are largely equivalent to wild-type in group 1 longevity mutants. *ife-2* mutants exhibit upregulation of *fmo-3, fmo-4* and *fmo-5*.

We also examined the expression of the other four *fmo* genes in the panel of long-lived mutants. In contrast to *fmo-2*, the expression of the four other *fmo* genes was largely unchanged in the group 1 longevity mutants (**Figure 1B-D**). There was a small but significant upregulation of *fmo-3, fmo-4* and *fmo-5* in *ife-2* mutants but the magnitude of this increase was smaller than the upregulation of *fmo-2* in the group 1 longevity mutants. Combined, this indicates that the upregulation of *fmo* genes in the group 1 longevity mutants is specific for *fmo-2*.

### *fmo-2* is required for lifespan extension in long-lived mitochondrial mutants

Having shown that *fmo-2* expression is significantly increased in the long-lived mitochondrial mutants, we next sought to determine if this increase in *fmo-2* might be contributing to their longevity. We also sought to confirm the increase in *fmo-2* expression using quantitative RT-PCR (qPCR). Similar to the RNA sequencing results, we found that *fmo-2* expression is significantly increased in *clk-1, isp-1* and *nuo-6* worms using qPCR (**Figure 2A-C**). While *fmo-2* RNAi had no effect on the lifespan of wild-type worms, knocking down *fmo-2* significantly decreased the lifespan of all three mitochondrial mutants *clk-1, isp-1* and *nuo-6* (**Figure 2A-C**). In each case, the lifespan following *fmo-2* RNAi was not fully reduced to wild-type lifespan. Thus, while *fmo-2* is required for the full lifespan extension in the long-lived mitochondrial mutants, other factors are also contributing to their longevity.

**Figure 2.**
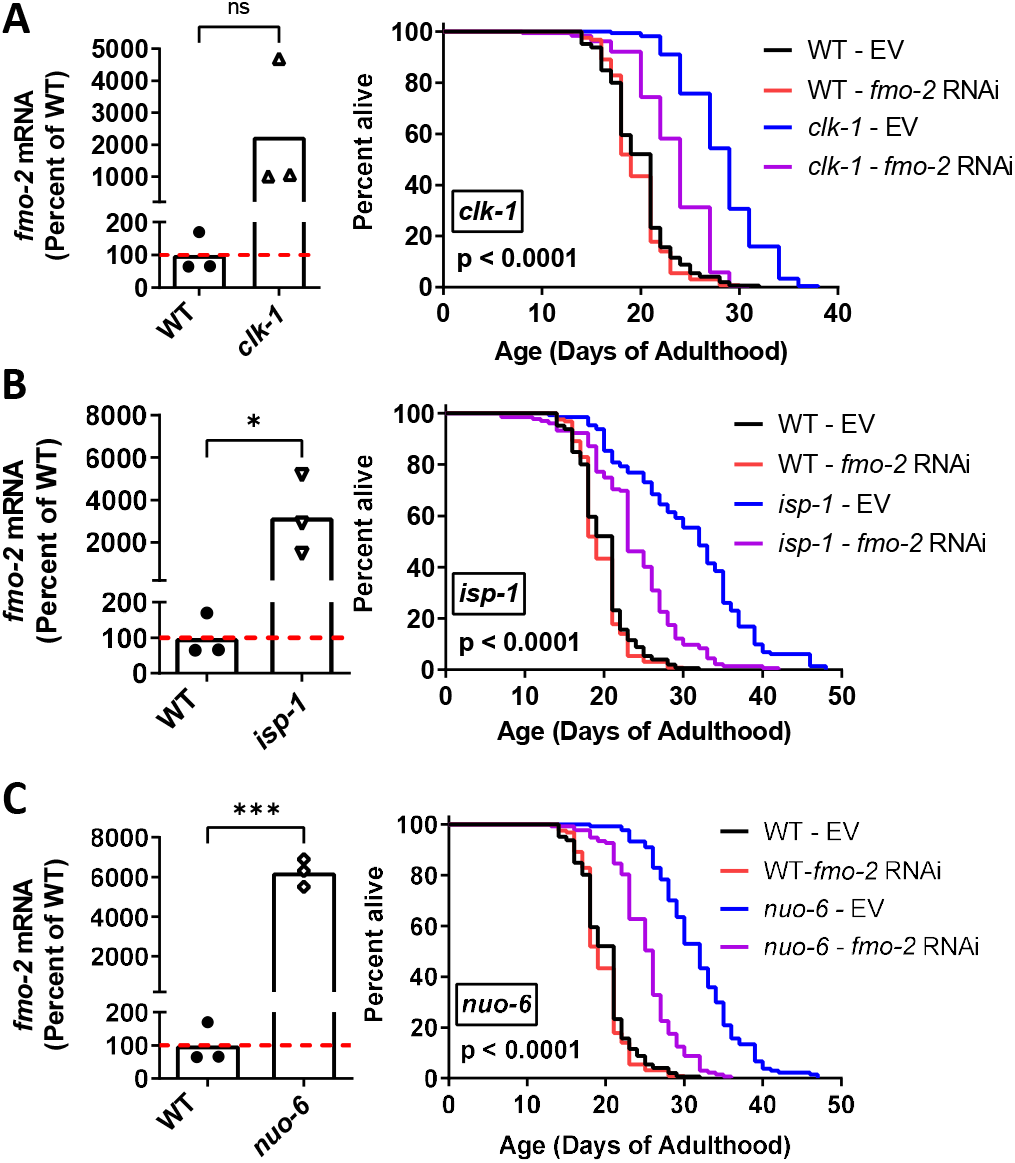
*fmo-2* is required for extended longevity in long-lived mitochondrial mutants. Quantification of *fmo-2* mRNA levels in day 1 pre-fertile young adult worms reveals that *fmo-2* mRNA levels are significantly increased in *clk-1* (**A**), *isp-1* (**B**) and *nuo-6* (**C**) worms. Decreasing *fmo-2* levels through treatment with *fmo-2* RNAi significantly decreased the lifespan of *clk-1* (**A**), *isp-1* (**B**) and *nuo-6* (**C**) worms despite having no effect on wild-type longevity. This suggests that the upregulation of *fmo-2* in these long-lived mutants is required for their extended longevity. Statistical significance was assessed using a student’s t-test for *fmo-2* mRNA levels or a log-rank test for survival plots. For survival plots, p value denotes the statistical significance between the blue and purple lines. Data for the effect of *fmo-2* RNAi on *nuo-6* lifespan is from Wu et al. 2018 *BMC Biology*.

To provide additional support for the role of *fmo-2* in the longevity of the long-lived mitochondrial mutants, we crossed the mitochondrial mutants to an *fmo-2* deletion mutant (ok2147) and measured lifespan. The ok2147 mutation is a 1070 bp deletion with a 2 bp insertion that affects exons 4 and 5 of 5 total exons. Consistent with the results from the *fmo-2* RNAi experiments, we found that deletion of *fmo-2* significantly decreased the lifespan of *clk-1* and *nuo-6* mutants (**Figure S1**). Note that we were not able to generate *isp-1;fmo-2* double mutants as both genes are in close proximity on the same chromosome. Combined, the *fmo-2* RNAi and *fmo-2* deletion results demonstrate that *fmo-2* is required for the long lifespan of the long-lived mitochondrial mutants.

### Transcription factors that activate *fmo-2* are required for longevity of long-lived mitochondrial mutants

Previous work has shown that the HLH-30 transcription factor is required for both the upregulation of *fmo-2* and lifespan extension resulting from HIF-1 activation, dietary restriction or *tald-1* knockdown ^5, 7^. The p38 MAPK PMK-1 was also shown to be required for *fmo-2* upregulation in response to dietary restriction or *tald-1* RNAi, and proposed to act in parallel with HLH-30 ^7^. Finally, it was found that MDT-15 and NHR-49 are required for the upregulation of *fmo-2* in response to organic peroxide, fasting or germline ablation (*glp-1* mutation), also in a HLH-30 independent manner ^8^.

Having shown that *fmo-2* is required for the longevity of the long-lived mitochondrial mutants, we sought to determine the extent to which genes upstream of *fmo-2* are also contributing to their long lifespans. To do this, we disrupted genes that were previously shown to be required for upregulation of *fmo-2* in the long-lived mitochondrial mutants and measured lifespan. We previously showed that disruption of genes involved in the p38-mediated innate immune signaling pathway (*nsy-1, sek-1, pmk-1*) are required for the long lifespan of *isp-1* and *nuo-6* worms ^25^. Knocking down *hlh-30* using RNAi significantly decreased the lifespan of *clk-1, isp-1* and *nuo-6* worms, but also reduced wild-type lifespan (**Figure 3A**). Similar to *fmo-2* knockdown, knockdown of *hlh-30* did not fully revert their lifespan back to wild-type treated with *hlh-30* RNAi. In contrast, we found that knocking down either *nhr-49* (**Figure 3B**) or *mdt-15* (**Figure 3C**) completely prevented lifespan extension resulting from mutations in *clk-1, isp-1* or *nuo-6* lifespan. Combined, this demonstrates that pathways involved in the activation of *fmo-2* are also required for the longevity of the long-lived mitochondrial mutants.

**Figure 3.**
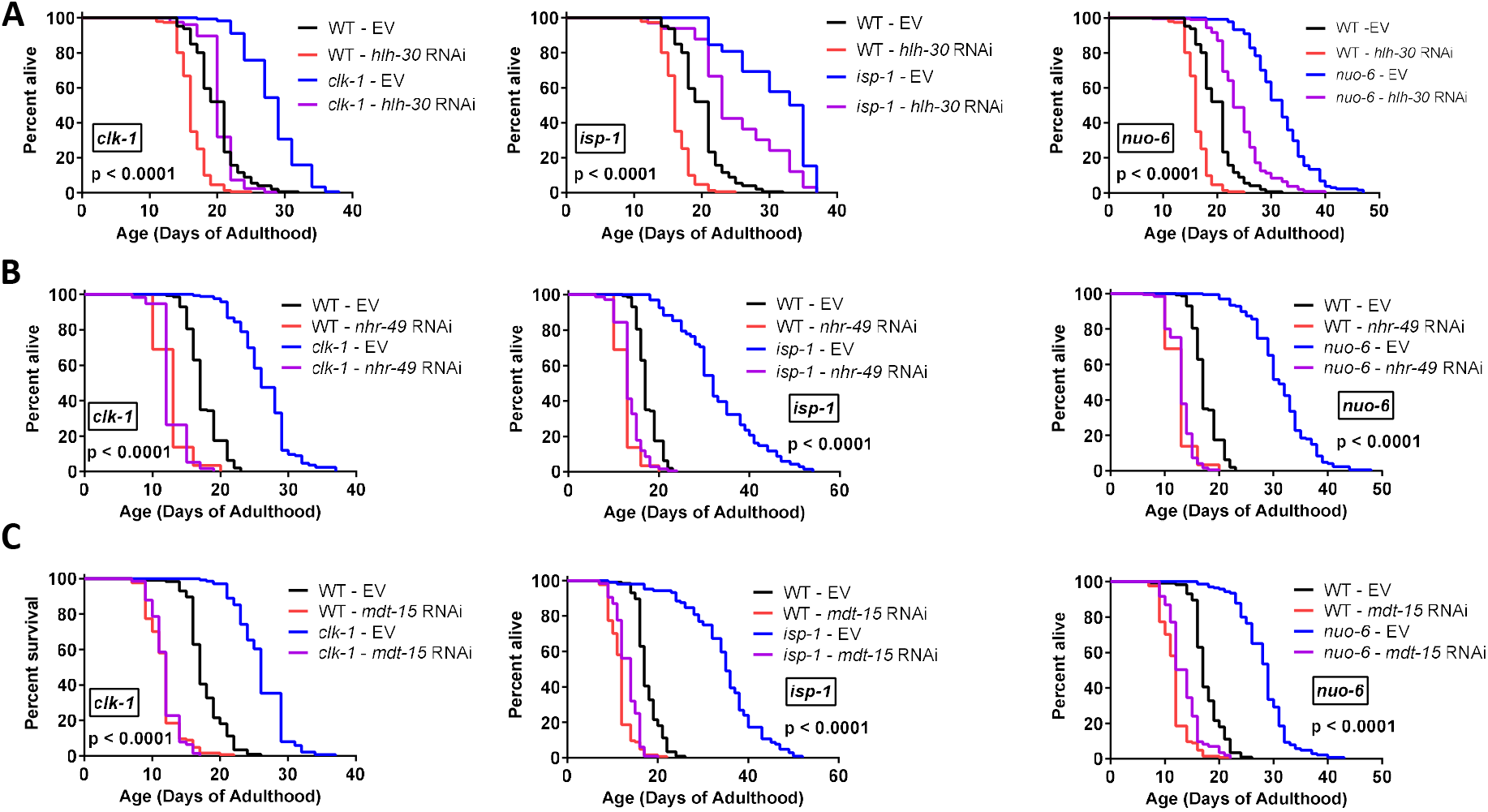
*hlh-30, nhr-49* and *mdt-15* are required for extended longevity in long-lived mitochondrial mutants and wild-type lifespan. Decreasing the expression of *hlh-30* (**A**), *nhr-49* (**B**) or *mdt-15* (**C**) decreased the lifespan of *clk-1, isp-1, nuo-6* and wild-type worms. The decrease in lifespan was partial for *hlh-30*, while the *clk-1, isp-1* and *nuo-6* mutations failed to increase lifespan when either *nhr-49* or *mdt-15* was knocked down. Statistical significance was assessed using a log-rank test. p value denotes the statistical significance between the blue and purple lines.

While these lifespan results are consistent with *hlh-30, nhr-49* and *mdt-15* contributing to the longevity of the long-lived mitochondrial mutants, it is possible that disruption of these genes is having a generally toxic effect as disruption of these genes also decreases lifespan in wild-type worms.

### Pathways that mediate lifespan of *clk-1* mitochondrial mutants contribute to upregulation of *fmo-2*

In a previous study, we observed that deletion of *atfs-1* decreased both *fmo-2* levels and lifespan in *nuo-6* mutants ^4^, suggesting that pathways affecting lifespan in the mitochondrial mutants strains may also be affecting *fmo-2* levels. To test this idea, we decided to focus on *clk-1* mutants. In exploring the molecular mechanisms of lifespan extension in *clk-1* mutants, we have identified a number of genes that are required for the long life of *clk-1* mutants, some of which have been previously reported by us or others. As our results suggest a role for *fmo-2* in *clk-1* longevity, we wondered which genes that affect *clk-1* lifespan also affect the expression of *fmo-2*. To study this, we disrupted specific genes related to longevity using either a genetic mutation or RNAi and then used quantitative RT-PCR to measure the levels of *fmo-2* mRNA in wild-type and *clk-1* animals compared to empty vector (EV) control. We also measured lifespan in all four groups.

We previously showed that disruption of the FOXO transcription factor DAF-16 or the p38 homolog PMK-1 both decrease *clk-1* lifespan ^13, 25^. Here, we extend these findings to show that *daf-16* RNAi (**Figure 4A**) or deletion of *pmk-1* (**Figure 4B**) both significantly decrease *fmo-2* levels in *clk-1* worms but not wild-type worms. The NRF2 homolog SKN-1 is involved in defending against oxidative stress and has been shown to be required for the long-lifespan of *daf-2* mutants ^26^. We find that in *clk-1* mutants, RNAi targeting *skn-1* decreases *clk-1* lifespan and reduces the expression of *fmo-2* (**Figure 4C**). Previous work showed that the nuclear homeobox protein CEH-23 is required for lifespan extension in *clk-1, isp-1* and *nuo-6* mutants ^27^. Deletion of *ceh-23* completely reverts the elevated *fmo-2* expression in *clk-1* worms back to wild-type but does not significantly affect *fmo-2* levels in wild-type worms (**Figure 4D**). The AMP-activated protein kinase AAK-2 has been shown to be required for the long lifespan of *daf-2* worms ^28^ and worms exposed to a lifespan-extending dose of paraquat ^29^. *Aak-2* is also partially required for the longevity of *clk-1* and *isp-1* mutants ^30^. In our hands, we find that deletion of *aak-2* reverts *clk-1* lifespan to wild-type and decreases the expression of *fmo-2* in *clk-1* worms (**Figure 4E**). Previous studies have shown that the hypoxia inducible factor HIF-1 is required for lifespan extension in *clk-1* and *isp-1* mutants ^3^. Here, we show that deletion of *hif-1* decreases both lifespan and *fmo-2* mRNA levels in *clk-1* worms (**Figure 4F**). The GATA transcription factor ELT-2 was shown to be required for wild-type lifespan and also the extended longevity of *daf-2, eat-2* and *isp-1* worms ^31^. RNAi targeting *elt-2* markedly decreases *clk-1* lifespan and reduces *fmo-2* expression specifically in *clk-1* mutants (**Figure 4G**). Combined, these results provide multiple examples of genetic pathways that are involved in both the longevity of *clk-1* worms and the expression of *fmo-2* in these mutants.

**Figure 4.**
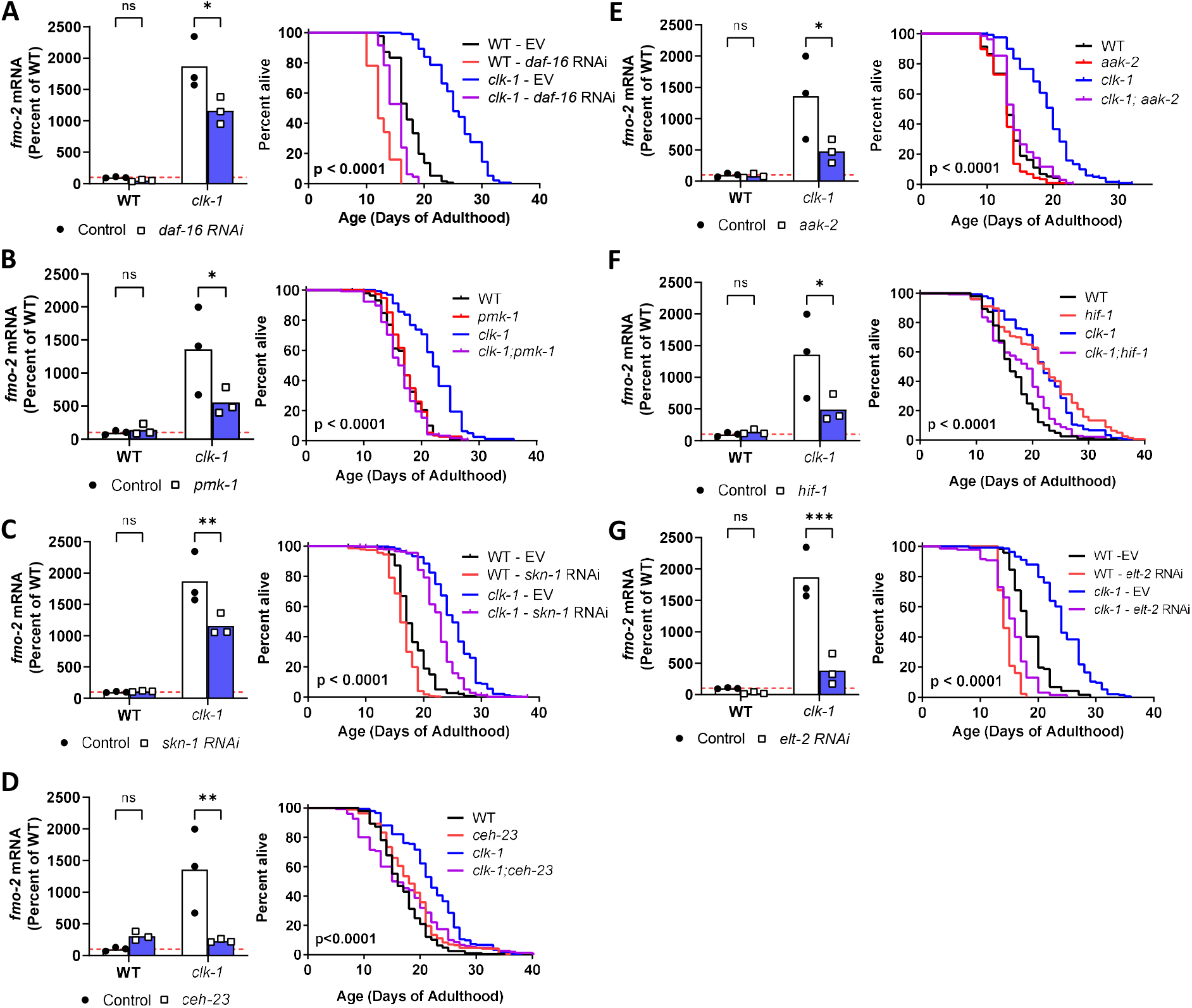
Pathways promoting longevity in long-lived *clk-1* mutants are specifically required for upregulation of *fmo-2* in *clk-1* worms. To better understand the relationship between *fmo-2* expression and lifespan in *clk-1* mutants, a number of lifespan-modulating pathways were disrupted in *clk-1* mutants using RNAi or genetic mutation before measuring lifespan and *fmo-2* expression. *fmo-2* expression was measured using qPCR in pre-fertile young adult worms. The genes that were disrupted included *daf-16* (**A**), *pmk-1* (**B**), *skn-1* (**C**), *ceh-23* (**D**), *aak-2* (**E**), *hif-1* (**F**) and *elt-2* (**G**). In every case, disruption of the longevity-related gene decreased *clk-1* lifespan and reduced *fmo-2* mRNA levels specifically in *clk-1* worms but not wild-type worms. Statistical significance was assessed using a two-way ANOVA with for Šidák’s multiple comparisons test for *fmo-2* mRNA levels or a log-rank test for survival plots. Lifespan data for DAF-16 is from Senchuk et al. 2018 *PLoS Genetics*. Lifespan data for PMK-1 is from Soo et al. 2023 *Aging Cell*. *p<0.05, **p<0.01, ***p<0.001.

## Discussion

In this work, we show that *fmo-2* is required for the extended longevity of the long-lived mitochondrial mutants, *clk-1, isp-1* and *nuo-6*. We also identify additional upstream pathways that contribute to *fmo-2* upregulation and provide further support for the positive relationship between *fmo-2* expression levels and lifespan. At the same time, we identify two long-lived mutants, *eat-2* and *osm-5*, that provide an exception to this correlation, as they have extended longevity despite reduced expression of *fmo-2*.

Previous studies have shown that *fmo-2* is upregulated by several factors including hypoxia, *vhl-1* deletion, dietary restriction, *tald-1* RNAi, *cco-1* RNAi, exposure to exopolysaccharide-producing strains of lactobacilli, exposure to tert-butyl hydroperoxide (tBOOH) and germline ablation (*glp-1* mutation) ^5, 7, 8^. Here, we extend these findings to show that *fmo-2* is upregulated in a specific group of longevity mutants that includes *clk-1, isp-1, nuo-6, daf-2, glp-1* and *sod-2* mutants. We previously showed that this group of long-lived mutants exhibits activation of the DAF-16-mediated stress response and the mitochondrial unfolded protein response ^18^.

Interestingly, long-lived mutants from a different longevity group, which includes *eat-2* and *osm-5* worms exhibit a significant downregulation of *fmo-2*. This provides a clear example in which *fmo-2* expression levels and lifespan extension can be experimentally dissociated. It suggests that these long-lived mutants are utilizing a different strategy for living long that does not rely on *fmo-2*.

Previous work has shown that the HIF-1 hypoxia pathway is activated in the long-lived mitochondrial mutants *clk-1, isp-1* and *nuo-6*, and that *hif-1* is required for lifespan extension in all three mutants ^3, 4^. Thus, it is plausible that *fmo-2* is activated in the mitochondrial mutants via the HIF-1 hypoxia pathway. This conclusion is supported by our observation that disruption of *hif-1* decreases *fmo-2* upregulation in *clk-1* worms. At the same time, it is likely that other pathways also contribute. In this work, we show that disruption of several signaling molecules that affect longevity are required for the upregulation of *fmo-2* in *clk-1* worms. In addition to *hif-1*, this includes *daf-2, pmk-1, skn-1, ceh-23, aak-2* and *elt-2*. Future epistasis experiments will be needed to sort out the extent to which these factors are working together or in parallel pathways to upregulated *fmo-2* expression.

Importantly, our data not only shows that *fmo-2* is upregulated in the long-lived mitochondrial mutants but also that *fmo-2* and factors involved in *fmo-2* upregulation are required for their extended longevity. In contrast to *isp-1* point mutants shown here to require *fmo-2* for their longevity, *fmo-2* is not required for *isp-1* RNAi to extend longevity ^5^. While on the surface these results are contradictory, it was previously shown that the mechanisms of lifespan extension in *isp-1* mutants and *isp-1* RNAi are different ^17^. This difference in *fmo-2-*dependency provides additional support for this conclusion.

Overall, this work highlights the role of *fmo-2* in determining longevity in the long-lived mitochondrial mutants. It also advances our understanding of the molecular mechanisms that modulate the expression of *fmo-2*. Based on these results, it appears that a mild impairment of mitochondrial function leads to the activation of a number of pathways that increase *fmo-2* expression and promote longevity. In future studies, it will be important to determine the extent to which these pathways act together or in parallel in promoting longevity and *fmo-2* activation.

## Conflict of Interest

*The authors declare that the research was conducted in the absence of any commercial or financial relationships that could be construed as a potential conflict of interest*.

## Author Contributions

Conceptualization: JVR. Methodology: JVR. Investigation: JVR. Visualization: JVR. Writing – original draft: JVR. Writing – review and editing: JVR.

## Funding

This work was supported by the Canadian Institutes of Health Research (CIHR; http://www.cihr-irsc.gc.ca/; JVR) and the Natural Sciences and Engineering Research Council of Canada (NSERC; https://www.nserc-crsng.gc.ca/index_eng.asp; JVR). JVR received a Senior Research Scholar career award from the Fonds de Recherche du Quebec Santé (FRQS) and Parkinson Quebec. The funders had no role in study design, data collection and analysis, decision to publish, or preparation of the manuscript.

## Acknowledgments

Some strains were provided by the CGC, which is funded by NIH Office of Research Infrastructure Programs (P30 OD010440). We would also like to acknowledge the *C. elegans* knockout consortium and the National Bioresource Project of Japan for providing strains used in this research.

---

**Figure S1.**
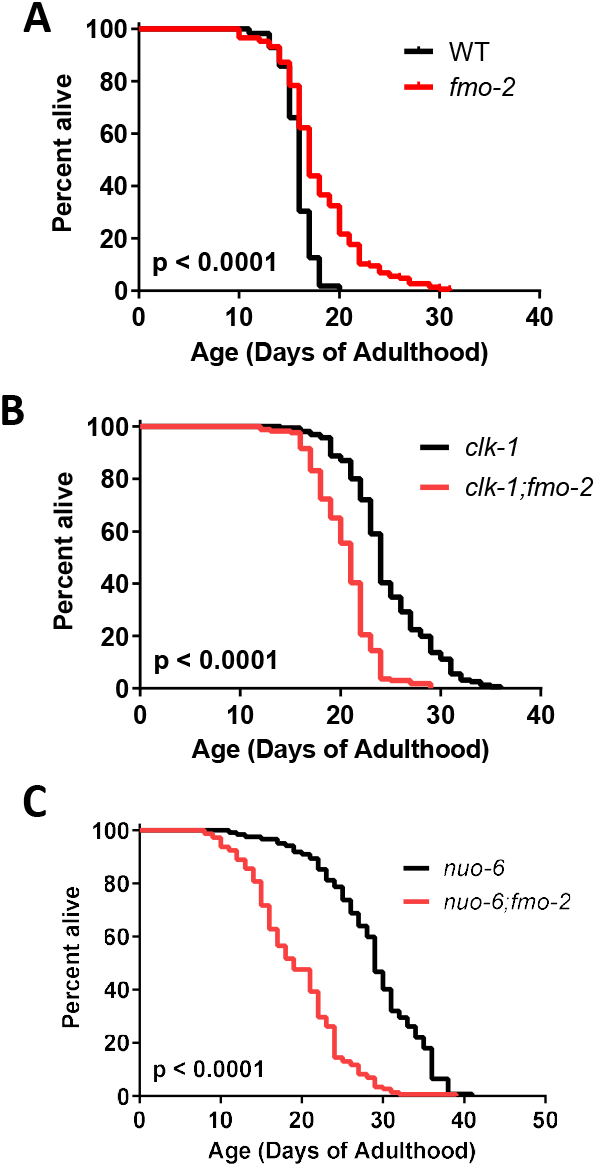
Deletion of *fmo-2* decreases the lifespan of long-lived mitochondrial mutants. While the *fmo-2* mutants exhibit increased lifespan in a wild-type background (**A**), deletion of *fmo-2* significantly reduces longevity in *clk-1* (**B**) and *nuo-6* (**C**) mutants. Statistical significance was assessed with a log-rank test.

## Notes

### Competing Interest Statement

The authors have declared no competing interest.

